# Host-associated phages disperse across the extraterrestrial analogue Antarctica

**DOI:** 10.1101/2021.11.09.467789

**Authors:** Janina Rahlff, Till L.V. Bornemann, Anna Lopatina, Konstantin Severinov, Alexander J. Probst

## Abstract

Extreme Antarctic conditions provide one of the closest analogues of extraterrestrial environments. Since air and snow samples, especially from polar regions, yield DNA amounts in the lower picogram range, binning of prokaryotic genomes is challenging and renders studying the dispersal of biological entities across these environments difficult. Here, we hypothesized that dispersal of host-associated bacteriophages (adsorbed, replicating or prophages) across the Antarctic continent can be tracked via their genetic signatures aiding our understanding of virus and host dispersal across long distances. Phage genome fragments (PGFs) reconstructed from surface snow metagenomes of three Antarctic stations were assigned to four host genomes, mainly Betaproteobacteria including *Ralstonia* spp. We reconstructed the complete genome of a temperate phage with near-complete alignment to a prophage in the reference genome of *Ralstonia pickettii* 12D. PGFs from different stations were related to each other at the genus level and matched similar hosts. Metagenomic read mapping and nucleotide polymorphism analysis revealed a wide dispersal of highly identical PGFs, 13 of which were detected in seawater from the Western Antarctic Peninsula in distance of 5538 km to the snow sampling stations. Our results suggest that host-associated phages, especially of *Ralstonia* sp., disperse over long distances despite harsh conditions of the Antarctic continent. Given that 14 phages associated with two *R. pickettii* draft genomes isolated from space equipment were identified, we conclude that *Ralstonia* phages are ideal mobile genetic elements to track dispersal and contamination in ecosystems relevant for Astrobiology.

**Importance:** Host-associated phages of the bacterium *Ralstonia* identified in snow samples can be used to track microbial dispersal over thousands of kilometers across the Antarctic continent, which functions as an extraterrestrial analogue because of its harsh environmental conditions. Due to presence of these bacteria carrying genome-integrated prophages on space-related equipment, and the here demonstrated potential for dispersal of host-associated phages, our work has implications for Planetary Protection, a discipline in Astrobiology interested in preventing contamination of celestial bodies with alien biomolecules or forms of life.

## Introduction

Due to harsh environmental conditions and isolation by the surrounding Southern Ocean’s Circumpolar Current, Antarctica is considered an analogue for multipurpose space exploration (1, 2). For example, its McMurdo Dry Valleys are regarded as a close terrestrial analogue to Mars (3). Astrobiology model organisms found on Antarctica are highly adapted to stressful conditions and comprise prokaryotes such as spore-forming *Bacilli* (4, 5), but also microfungi (3, 6). Understanding endurance and dispersal of microorganisms under conditions that mimic those on extraterrestrial planets, i.e., high UV radiation, low temperature, and low nutrient availability, has important implications for Planetary Protection. For instance, the dispersal of microbes that hitchhike to a celestial body is currently not considered in Planetary Protection, a discipline in Astrobiology set out with the aim of preventing contamination of celestial bodies with foreign biomolecules or forms of life.

Among the potential candidates hitchhiking spacecraft are Betaproteobacteria of the genus *Ralstonia* (order *Burkholderiales*). These bacteria are able to thrive under oligotrophic conditions (7) and were reported to be ubiquitously present on space-related equipment including water systems of the International Space Station (ISS) (8, 9), and the Mir space station (10, 11). Likewise, they belonged to the microbial inventory of Mars Odyssey and Mars Phoenix lander facilities and can thus prevail under strict Planetary Protection regulations (12, 13). *Ralstonia pickettii* strains were found to thrive in simulated microgravity compared to normal gravity (11) and demonstrated high resistance against different metal ions and UV-C radiation (8).

*Ralstonia* spp., (mainly *R. pickettii*), were previously found Antarctic soils (14), Antarctic snow (15–17), in snow over Tibetan Plateau Glaciers (18), and in the air of the Antarctic base Concordia (19). Interestingly, this genus has been regarded as an atmospheric traveler rather than being part of true snow microflora in Antarctica (15). Despite a report on bacterial activity at subzero temperatures in South pole snow (20), *Ralstonia* was not among active bacterial communities as inferred from cDNA-based 16S rRNA amplicon sequencing (15). This view was supported by a comprehensive Antarctic surface snow microbiome study that did not detect this bacterial genus (21), and by the fact that spatial variability of snow microbiomes in Antarctica is high (22). Although aerial dispersal is probably the major contributor to (micro-)biological input to remote regions (23), the role of bioaerosol transport to microbial ecology of isolated systems such as the Antarctic continent is poorly understood (24). Applying high throughput sequencing approaches to study bacterial dispersal over the Antarctic continent remains a considerable challenge due to the low microbial biomass of atmosphere-derived samples regarding their DNA content (25), and resulting issues of recovering high quality assemblies from metagenomic reads (26).

In addition to the limited knowledge about how transport via aerosols and snow across Antarctica shapes microbial dispersal patterns, another open question relates to the distances that microbes can cover within the atmosphere of extreme environments. A study on the dispersal of airborne faecal coliforms showed a distribution over about just 175 m from a sewage outfall at Rothera Research Station (Antarctic Peninsula) and thus prolonged survival was considered unlikely (27). More stress-resistant microorganisms could, however, endure for much longer periods. Recently, L. A. Malard et al. (22) found high abundances of spore-forming *Bacilli* and suggested that long-term dispersal may seed continental Antarctic snow ecosystems. However, to date, the role of aerial dispersal in shaping patterns of microbial biogeography is supported by little empirical evidence (23).

Here, we follow the hypothesis that geographically widespread (28) *Ralstonia* spp. prophages and/or replicating and adsorbed bacteriophages of this genus (all types further referred to as “host-associated” phages) can be used to study host bacterium dispersal across the Antarctic continent. Reconstructing prokaryotic genomes from samples containing low amounts of DNA, including air or precipitation is challenging, as low input libraries (∼1 pg) can result in problems of genome binning (26). However, (pro)phage genomes or their fragments are much smaller than prokaryotic genomes and thus easier to identify, track and compare. In this study, genome-resolved metagenomics was applied to demonstrate dispersal of host-associated phage genome fragments (PGFs) from surface snow across the Antarctic continent over hundreds to thousands of kilometers. We detected PGFs belonging to the orders *Caudovirales* and *Tubulavirales* in this extraterrestrial analogue and additionally show that similar genome-integrated phages are frequent colonizers of space equipment. Therefore, we suggest that host-associated PGFs represent a useful tool to study spatial dispersal of bacteria and their phages in extreme environmental settings and further envision implications for the dispersal of microbiological contaminations on spacecraft and celestial bodies that previously escaped Planetary Protection measures.

## Results

### Reconstruction of low coverage MAGs from Antarctic snow metagenomic data

The microbial community composition of surface snow samples collected close to three Russian Antarctic stations, Druzhnaja, Mirnii and Progress, based on 16S rRNA gene sequencing was described earlier (16) and showed that *Ralstonia* was the most dominant organism in snow collected at the Mirnii station, the second most dominant after *Janthinobacterium* at Druzhnaja, and the third most dominant (after *Flavobacteria* and *Hydrogenophaga*) around Progress station. We reconstructed four prokaryotic and one eukaryotic metagenome assembled genomes (MAG) from the three low-biomass snow metagenomes (Figure 1A). Two MAGs related to *Ralstonia* sp. and *R. pickettii* were recovered from Druzhnaja and Mirnii samples and had 86%/10% and 55%/0% completeness/contamination scores, respectively. MAGs of *Janthinobacterium lividum* with 92%/6% scores and of *Flavobacterium micromati* with 96%/0% scores were recovered from Druzhnaja and Progress, respectively. We also identified a MAG of a diatom (likely *Thalassiosira* sp.) in the Mirnii metagenome, which we used for normalizing some of our viral analysis (see below) but was not further characterized in this work. Read mapping revealed coverage scores of 5.9, 7.1, 6.7 and 14.6 for the MAGs of *Ralstonia* sp.*, R. pickettii, J. lividum* and *F. micromati*, respectively.

**Figure 1:**
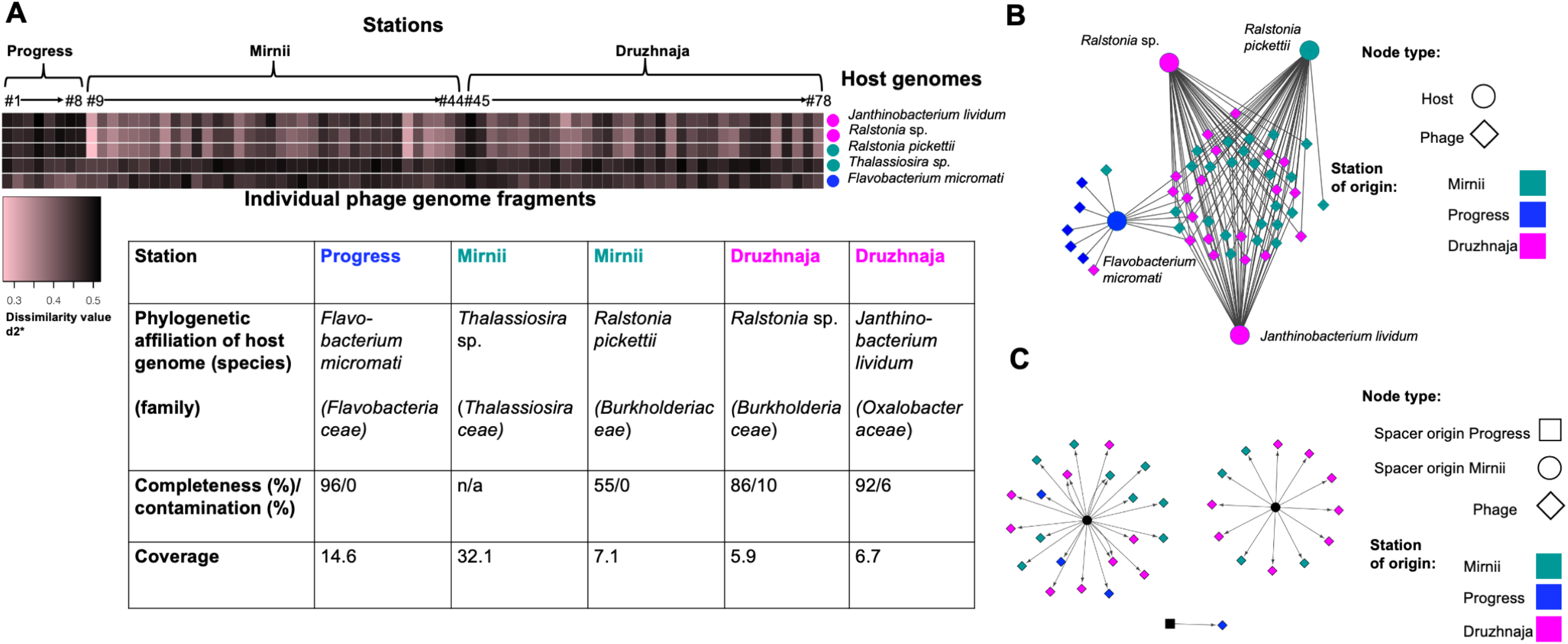
Host-phage pairings based on shared k-mer frequency patterns. **A)** Heatmap representing dissimilarity value d2* for matches between five host MAGs and 78 PGFs derived from three Antarctic snow metagenomes from the stations Progress, Mirnii, and Druzhnaja. The PGF number corresponds to the number in the phage name, for instance PGF#78 refers to Antarcphage78_Dr_3477 (Table S1). The table shows MAG characteristics (phylogenetic affiliation, completeness, contamination, and coverage). n/a = not available **B)** Network of phage-host interactions based on k-mer frequency pattern reveals strong overlap between *Ralstonia* and *Janthinobacterium lividum* infecting PGFs. Here, only PGFs matching the host MAG below the defined threshold (see text) are shown. **C)** CRISPR spacer matches to PGFs. Different spacers matching to the same PGFs are shown by multiple arrows. The two hosts of CRISPR arrays for Mirnii-derived spacers remain unidentified according to their direct repeat sequence, whereas the Progress spacer is derived from a *Flavobacterium* sp. (Table S2).

### Prevalent absence of CRISPR-Cas systems suggest low adaptive immunity

We searched for clustered regularly interspaced short palindromic repeats (CRISPR) spacers in the snow metagenome reads to link them to potential protospacers on PGFs. CRISPR arrays and *cas* genes were absent from both *Ralstonia* and the *J. lividum* MAGs as determined by CRISPRcasFinder (29). BLASTing of direct repeat (DR) sequences to NCBI’s NR database also did not indicate *Ralstonia* or *Janthinobacterium* to be the host of the CRISPR array. Solely the genome of *F. micromati* contained two CRISPR arrays with four spacers each (both evidence level 3 = highly likely candidates). CRISPR arrays were not detected on *R. pickettii* draft genomes SSH4 and CW2 obtained from space equipment.

### PGF-host analyses suggest shared hosts of Antarctic phages

A total of 26 predicted PGFs, eight of them being putative PGFs, was found in Antarctic snow metagenomic data (Table S1). VIBRANT (30) additionally detected 52 PGFs, resulting in a total of 78 PGFs (Figure 1A). PGFs with minimum 75% of the genome covered with reads had coverages ranging between 1.9 and 17.5. Most PGFs were partial sequences according to viralComplete (31) and CheckV (32), only Antarcphage10_Mi_4716 and Antarcphage49_Dr_7823_circ were estimated to be of full length (Table S1), although annotations of Antarcphage10_Mi_4716 in comparison to *Ralstonia* phage p12J revealed missing genes and let us question its completeness (Figure 2). Out of 78 PGFs, 77 were categorized as lytic by VirSorter (33) and 75 were found to be of viral or unclassified origin by CheckV (Table S1). A proportion of 43.6%, 46.1% and 10.3% identified PGFs originated from the Druzhnaja, Mirnii and Progress, respectively.

**Figure 2:**
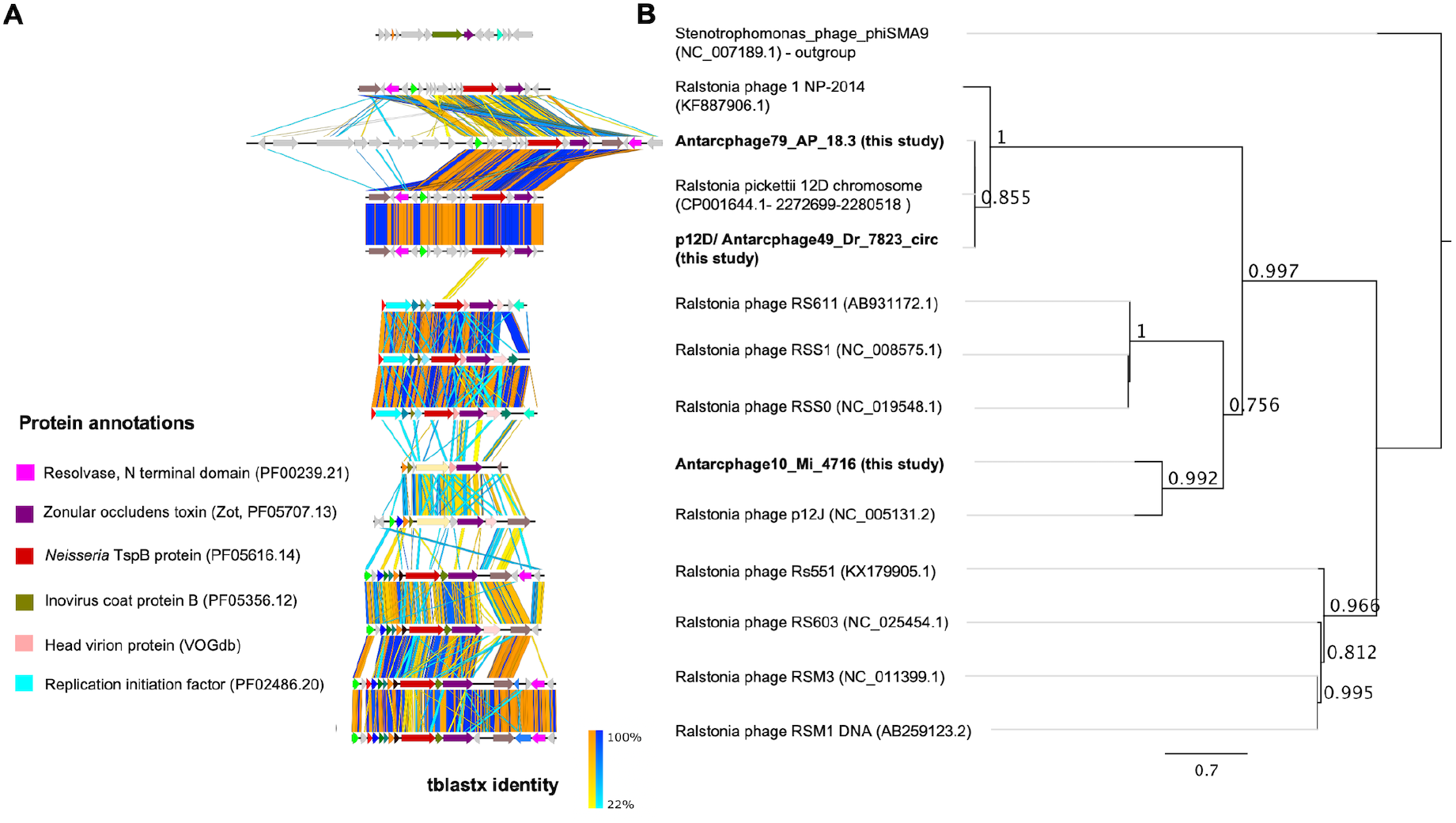
Phage genome comparisons, functional annotations and phylogenetic relationship based on zonular occludes toxin (Zot, Pfam-ID: PF05707). **A)** Synteny of known *Ralstonia* sp. phages from NCBI and Antarctic PGFs from this study with similar coding sequences (CDS) being in the same color if they occur on more than one phage genome. Functional annotations performed by DRAM-v (115) are given for all colored CDS where available. Vertical lines between sequences indicate regions of shared similarity shaded according to tBLASTx (orange gradient for matches in the same direction or blue gradient for inverted matches). Figure was created using Easyfig (119). **B)** The phylogenetic tree was built using FastTree 2.1.11 (122) in Geneious 11.1.5 (112) under default settings and shows four distinct clusters of Zot proteins based on their amino acid sequences (aligned with MUSCLE (121)).

All 78 PGFs were analyzed in conjunction with reconstructed MAGs using VirHostMatcher (34). We defined a dissimilarity threshold of d2* value = 0.436, which corresponds to the lowest dissimilarity value (= highest similarity) for PGFs matching the MAG of *Thalassiosira* sp. from the snow environment (Figure 1A) by assuming that the eukaryotic MAG does not match any of our extracted prokaryote infecting PGFs. In total, 50 of the 78 PGFs matched a host MAG below the defined dissimilarity threshold, based on their shared k-mer patterns. Most PGFs (41) matched both MAGs of *Ralstonia* sp. and *R. pickettii* (Figure 1A). In total, 38 and 14 PGFs matched *J. lividum* and *F. micromati* MAGs, respectively. Five out of eight PGFs extracted from the Progress station metagenomic snow sequences matched the *F. micromati* MAG recovered from this station. Seven PGF matches were shared between all prokaryotic MAGs (Figure S1). However, with 30 shared matches, the three MAGs belonging to the order *Burkholderiales* shared the most overlap (Figure 1B, Figure S1).

Matching of CRISPR spacers derived from CRISPR loci of unknown hosts (reconstructed from DR sequences) revealed that spacers from Mirnii matched 19, 14 and three PGFs from Druzhnaja, Mirnii and Progress, respectively, and one *Flavobacterium* sp. spacer from Progress matched a PGF from that station (Figure 1C, Table S2, ≥80% similarity). Out of these 37 spacer matches, three matched protospacers of PGFs of unknown hosts, and 26 were assigned to a *Ralstonia* host according to VirHostMatcher or other predictions (Table S1). Since p12D (see below) was among the PGF spacer targets and is a certain *Ralstonia* phage, this could indicate that an unknown host belonging to the order *Burkholderiales* uses adaptive immunity against viruses. We compiled evidence of host prediction for the 78 PGFs in Table S1.

### The temperate phage p12D forms a distinct, monophyletic clade excluding most known *Ralstonia* phages

Among the 78 PGFs, we detected a circular (and thus complete) 7.8 kb PGF termed Antarcphage49_Dr_7823_circ (Figure 2A, Figure S2). Since the Antarcphage49_Dr_7823_circ PGF from the Druzhnaja station was found in the chromosome of a *R. pickettii* 12D strain isolated from copper-contaminated sediment from a lake in Michigan with 99.9% identity (accession number: CP001644.1-2272699-2280518, Figure 2A), we here propose the name p12D phage equivalent to *Ralstonia* p12J, a known phage infecting the *R. pickettii* 12J strain. Antarcphage49_Dr_7823_circ (from now on referred to as p12D) only matched the MAG of *Ralstonia* sp. and *R. pickettii* based on shared k-mer frequency patterns. Functional protein annotations provided evidence that p12D contained a gene for a resolvase domain containing protein/site-specific recombinase, likely used for integration into the host genome, as well as for the zonular occludens toxin (Zot, PF05707) (Figure 2A, Figure S2, Table S3). The non-toxic component of Zot at the N-terminus represents a characteristic protein in filamentous phages that has been used for phage classification (35, 36). Reconstruction of a phylogenetic relationships of Zot proteins from this study and respective references (Figure 2B, Figure S3) confirmed a close identity of p12D to *Ralstonia* phage 1 NP-2014 and Antarcphage79_WAP_18.3, all belonging to the same monophyletic clade. *Zot* of Antarcphage10_Mi_4716 was phylogenetically related to filamentous *Ralstonia* phage p12J. The tree shows four distinctive clusters for the Zot protein, reflecting the overall synteny of the gene order of the different *Ralstonia* phages very well.

Annotations against UniRef100 revealed that 68.8% of all annotated Antarctic PGF proteins remain hypothetical. The annotations taxonomically assigned 34.4% of all Antarctic PGF proteins to either *Ralstonia* or *R. pickettii,* and 11.6 % to phages of *Janthinobacterium* or *J. lividum*, potentially indicating lateral gene transfer between virus and host in their respective evolution (37, 38) or supporting that these PGF indeed represent prophages. Three of the Antarctic PGFs carried a site-specific integrase or resolvase domain, and 4.2% of genes were related to phage structural proteins, e.g., head, tail, or capsid proteins (Table S4, Table S5).

### Cross-mapping and nucleotide variations reveal dispersal patterns of MAGs and PGFs across Antarctica

Mapping of 100% identical reads from the three snow samples and the Western Antarctic Peninsula (WAP, Figure 3A) seawater sample to the prokaryotic MAGs revealed that both *Ralstonia* MAGs were detected at all sites (89-100% genome coverage). *J. lividum* was considered absent from the Progress station (genome coverage of 62%) and detected at other stations (minimum 95% covered genome). *F. micromati* was only detected in Progress (100%) and Druzhnaja (96%) but considered absent from Mirnii (58.7%) and the WAP (1.8%). Cross-mapping on the 78 PGFs demonstrated that PGFs derived from the Progress station could not be found in the other two snow metagenomes (Figure 3B&C). By contrast, 16 different PGFs from Druzhnaja were detected at Mirnii, and 27 Mirnii PGFs were found at Druzhnaja (Figure 3C). In addition, 4 and 9 PGFs from Mirnii and Druzhnaja were detected in the WAP dataset, respectively (Figure 3 B&C, all of them having minimum 97% of their lengths covered with reads from the snow metagenomes (Table S1). Of 43 PGFs that were found at both stations (Mirnii and Druzhnaja) based on read mapping, 26 PGFs shared a viral cluster (VC), and 34 matched prokaryotic hosts based on k-mer frequencies between stations (Table S1). Interestingly, nine of the 13 PGFs occurring in WAP and Mirnii/Druzhnaja samples contained identical single nucleotide polymorphisms (SNPs, Figure 3D&E, Table S6) or were missing common SNPs pointing towards a common phage population before dispersal led to separation.

**Figure 3:**
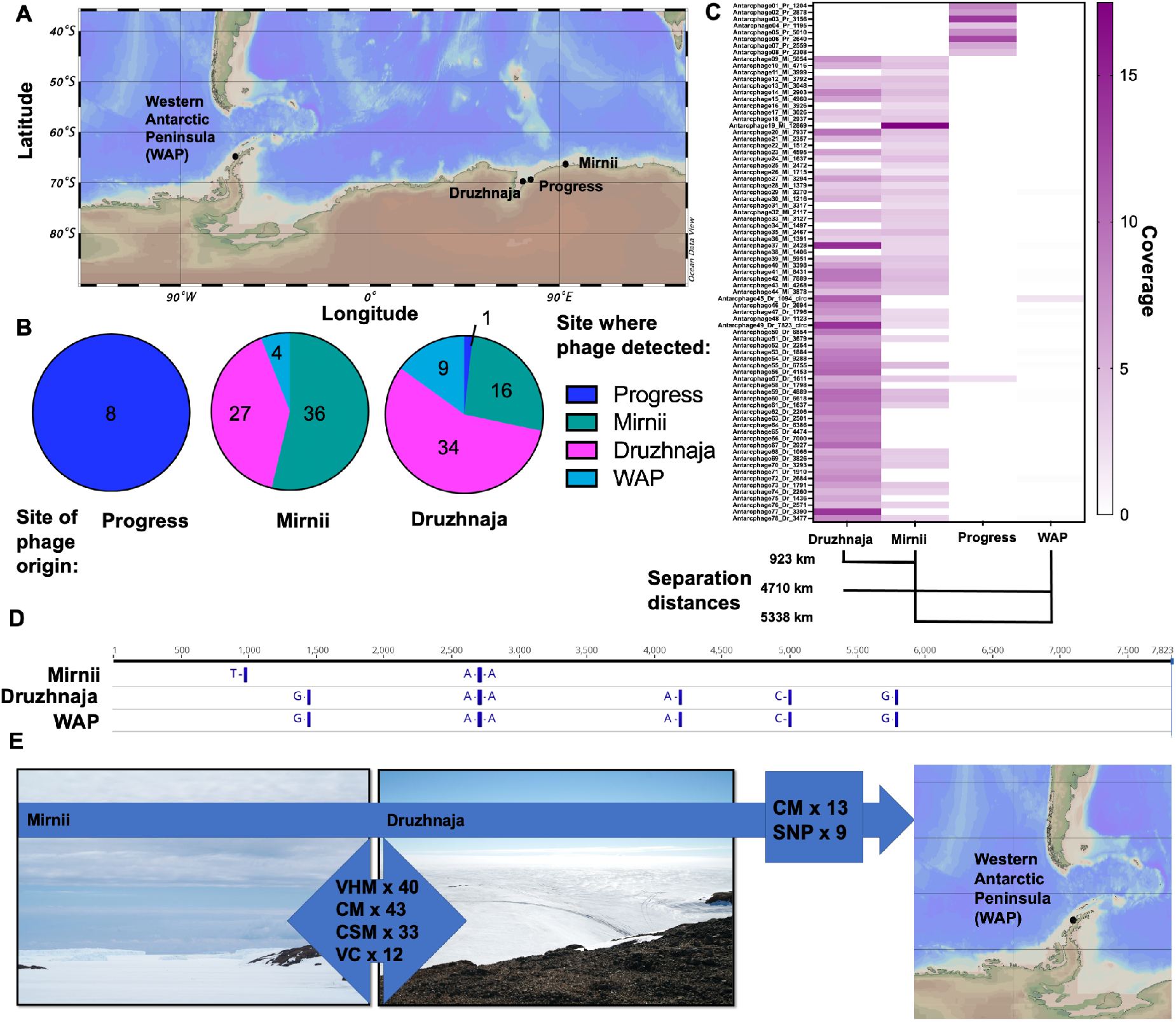
Evidence for dispersal of PGFs across the Antarctic continent. **A)** A map showing the different snow sampling stations in the East and the sampling station for seawater at the Western Antarctic Peninsula. Map was built using Ocean Data View (123). **B)** Pie charts summarizing the number of PGFs considered present at each station based on cross mapping of reads and the site of PGF assembly (phage origin). **C)** The heat map depicts the normalized coverages based on cross-mapping of reads against the 78 PGFs and separation distances between stations. White areas indicate PGFs that were not called present because they had less than 75% of scaffold length covered in mapping (with least 1-fold coverage). **D)** Single-nucleotide polymorphism (SNP) analysis of p12D/Antarcphage49_Dr_7823_circ based on reads from Druzhnaja, Mirnii and the WAP. This PGF was absent from the metagenome of the Progress station. Further SNP analysis data can be found in Table S6. **E)** Summary figure for the major dispersal route and supporting evidence. The majority of PGF disperse between Mirnii and Druzhnaja, and 13 PGFs additionally occurred at the WAP based on cross-mapping (CM) and SNP analysis. Values refer to the number of tested features, which include number of virus clusters (VC), number of shared viruses based on CM, virus-host matches based on k-mer links (VHM), CRISPR-spacer matches (CSM). A VHM=1 indicates that one PGF has infected host MAGs from two stations. No VC and only one CM were observed for Progress-Druzhnaja and Progress-Mirnii. Therefore, this station was excluded from the summary.

Clustering of PGFs in VICTOR (39) and vConTACT2 (40) revealed that groups of two to four PGFs could be assigned to twelve distinct viral genera-forming viral clusters (Figure 3E, Figure S4 & S5, Table S1). Members of the same cluster were often recovered from different stations, for example, Druzhnaja and Mirnii (Figure 3B), which are located 923 km apart (Figure 3C). The phylogenomic Genome-BLAST Distance Phylogeny (GBDP) tree yielded average support of 47% and 80% in the nucleic acid and amino acid-based analysis, respectively. OPTSIL clustering yielded 78 and 64 species and genus clusters at the nucleic acid level and 78 and 70 at the amino acid level, respectively.

Many PGFs formed VCs with no genomic relatedness to any known phages from the ProkaryoticViralRefSeq v94 database. Others were, however, similar to known *Ralstonia* PGFs (current family *Inoviridae,* order *Tubulavirales*) and hence clustered with those, such as p12D with *Ralstonia* phage 1 NP-2014, or Antarcphage10_Mi_4716 with *Ralstonia* phage PE226 and *Ralstonia* phage p12J (Figure 4, Table S1). Other Antarctic PGFs shared protein clusters with *Ralstonia* phages from space equipment based on vConTACT2 or were related to phages of the *Caudovirales* order from the RefSeq database (Figure 4).

**Figure 4:**
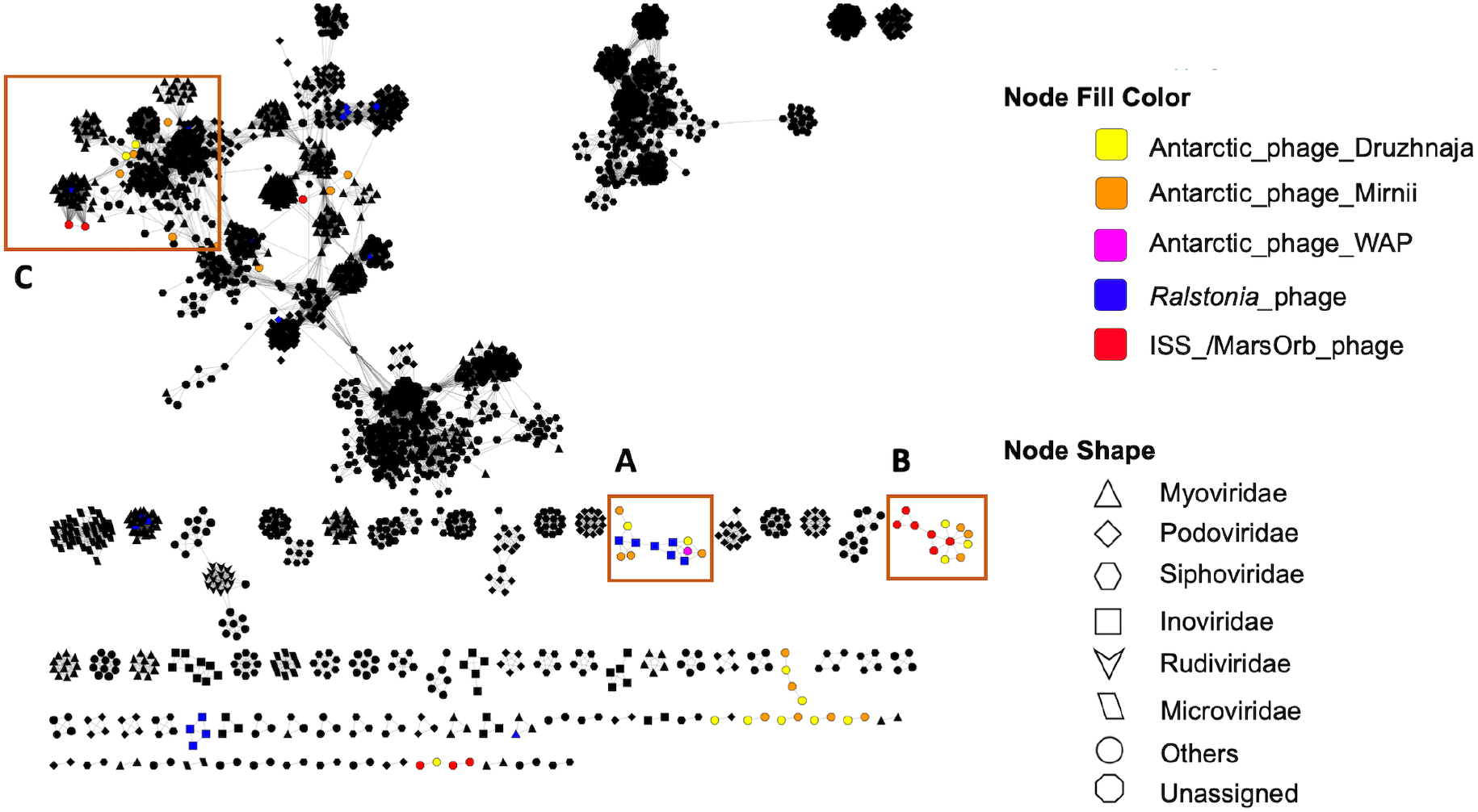
Phage network of Antarctic PGFs derived from Druzhnaja, Mirnii, Western Antarctic Peninsula (WAP) and space equipment (ISS/MarsOrb) clustered with viruses of the viral RefSeq database. Based on shared protein clusters, Antarctic PGFs from this study group with known *Ralstonia* phage of the family *Inoviridae* (A), with phages obtained from *Ralstonia* isolates from space equipment (B) or with known phages of the order *Caudovirales* (C). Black nodes refer prokaryotic viruses other than *Ralstonia* and PGFs from this study. Interactions show relatedness of genomes on viral genus or higher taxonomic level. “Others” refer to other known viral families not listed in the legend. Visualization was done using Cytoscape 3.8.2. (109). For details about viral clusters, please see Table S1.

In summary, our results support the idea of dispersal of host-associated phages between widely separated stations because a) cross-mapping revealed presence of Mirnii PGFs at Druzhnaja and vice versa, presence of a temperate phage from snow in the WAP seawater dataset, and presence of the hosts *Ralstonia* and *J. lividum* in metagenomes of most stations; b) PGFs in widely separated locations carry identical SNPs or lack nucleotide variations; c) some PGFs from Mirnii and Druzhnaja belong to the same genus/VC; and d) Mirnii and Druzhnaja PGFs share host MAGs according to their k-mer frequency patterns. These observations were mainly true for PGFs affiliated to *Ralstonia* and *J. lividum* MAGs and less so for *F. micromati* PGFs, which could indicate that certain bacterial species or their phages in Antarctic snow are more prone to atmospheric dispersal than others. Finally, we found CRISPR spacers from two unidentified hosts (represented by two distinct CRISPR DR sequences) at the Mirnii station to match PGFs from Druzhnaja, Mirnii and Progress. Mapping revealed little to no abundance of Progress PGFs at other stations, and no *F. micromati* MAG was detected in Mirnii and WAP samples. This could be related to the low biomass and/or or insufficient sequencing coverage. However, spacers from Mirnii suggest an infection history with PGF from Progress, thus reflecting (past) dispersal of host and/or phage.

### *Ralstonia* phages occur across diverse ecosystems and on space equipment

By BLASTing all recovered PGFs against the IMG/VR viral database (41), we found that p12D shared high identity (85.9 %) with a *R. pickettii* prophage (IMG/VR v.2 scaffold ID Ga0075447_10000781, genome ID: 3300006191) from a seawater metagenome of the WAP (Figure 2), which was further validated by cross-mapping of WAP reads to p12D delivering 100% scaffold coverage with 90% identical reads. The WAP is 4710 km away from the Druzhnaja station (Figure 3C). The PGF Antarcphage79_WAP_18.3 was obtained by BLASTing the p12D scaffold against the WAP assembly but was assembled containing genomic regions extending the actual PGF. Based on vConTACT2, it was affiliated to the same viral genus as p12D. Other PGFs recovered from the Antarctic snow metagenome showed hits to phages deposited at IMG/VR. Hits were related to phages originating from freshwater, wastewater, groundwater, or phages that were associated with plant root microbial communities, which is in accordance with the fact that *Ralstonia* sp. is a frequent phytopathogen (42). Many of the matching entries in IMG/VR were identified as phages of *Ralstonia* due to matching with CRISPR spacers (mostly of *R. solanacearum,* Table S7).

Altogether, 14 PGFs were found on space equipment, hereafter referred to as ISSphage (7) and MarsOrbphage (7), for PGFs extracted from the *R. pickettii* strains CW2 and SSH4 draft genomes from the ISS cooling system and Mars Odyssey Orbiter, respectively. Seven of these PGFs were identified as lysogenic and according to protein annotations (Table S4 & S5) rely on proteins of the integrase family for integration into the host chromosome. vConTACT2 revealed that some of the ISSphage4 formed a common genus cluster with four Antarctic PGFs, MarsOrbphage5 shared protein clusters with ISSphage3, and MarsOrbphage7 formed a genus cluster with several *Burkholderia* phages among other *Caudovirales* members (Figure 4, Table S1).

## Discussion

CRISPR-Cas systems equip bacteria and archaea with a powerful defense system against invading mobile genetic elements including plasmids, phages and viruses. Only ca. 50% of bacteria rely on this adaptive immune system, and only 31% of genomes from public databases belonging to plant-pathogenic *Ralstonia solanacearum* contain CRISPR-Cas arrays (43). Upon exposure to a virulent phage under laboratory conditions, the CRISPR array of *R. solanacearum* strain CFBP2957 did not acquire new spacers from viral protospacers (43). O. S. Gonçalves et al. (28) reported that in the presence of CRISPR arrays, 27.9% of CRISPR spacers from *Ralstonia* genomes targeted prophage elements. In our study, two *Ralstonia* MAGs from a low temperature environment were devoid of CRISPR systems, implying that representatives of this genus have alternative strategies for defense against mobile genetic elements (43). However, analyses on more *Ralstonia* genomes are necessary to corroborate this statement.

Based on shared protein clusters, some Antarctic PGFs such as Antarcphage10_Mi_4716, Antarcphage48_Dr_1123, Antarcphage17_Mi_3026, and p12D clustered with known *Ralstonia* phages including *Ralstonia* phage p12J (NC_005131.2), *Ralstonia* phage PE226 (NC_015297.1) or *Ralstonia* phage 1 NP-2014 (NC_023586.1). These are filamentous phages of the *Inoviridae*, a family that has been recently called for reclassification to higher taxonomic ranks (36, 44). Inoviruses (order *Tubulavirales*) typically feature circular, single stranded DNA genomes of ∼5-15 kb length, lead to chronic infections, and are globally abundant (36). As our protocol should have only revealed double-stranded DNA, detection of ssDNA inoviruses sequences shows that these must be either replicating phages or genome-integrated prophages. *Zot*, which was detected on p12D, Antarcphage10_Mi_4716 and Antarcphage79_WAP_18.3 is a typical gene in filamentous *Ralstonia* phages (45, 46). This phage type confers little burden to its host or can even serve it, e.g., by increasing its hosts’ virulence and evolutionary fitness, and because virions can leave the host in a non-destructible way (35, 36). *Ralstonia* phages detected in Arctic viromes were shown to transduce genomic information of cold-shock proteins to their hosts (47), a clear asset for microorganisms in polar environments. However, we did not find transduction of beneficial genes, which seems to be common in extreme environments (48).

Some *Ralstonia* phages occur as non-integrative, episomal forms, e.g., RS603, a hybrid of RSM1/3 infecting the phytopathogen *R. solanacearum* (49), whose genome lacks a resolvase domain (Figure 2A), but many mesophilic *Ralstonia* also occur as lysogens (50). Lysogeny, a lifestyle during which the phage genome becomes integrated into the host chromosome, is a widespread phenomenon in low temperature environments (51–53) and likely attributed to prolonged starvation and low activity of host cells under harsh conditions, the latter being previously reported for *Ralstonia* (15). Since p12D and its counterpart from the WAP have a resolvase-domain containing protein likely functioning in integration/excision during lysogenisation (54, 55), we conclude that they must be temperate phages of *Ralstonia*.

Many, i.e., 75 of the 78 Antarctic PGFs found in this study shared little protein clusters with known phages from public databases. This is certainly related to the often high diversity of viruses, limited accessibility to Antarctic environments as well as a stronger focus on sequencing metaviromes and viral isolates of direct human interest (56). The missing relatedness is known for ssDNA viruses originating from Antarctic cryoconite holes and was attributed to the isolation and extreme environmental conditions at the Antarctic continent (57). In total, 45 of the 78 PGFs could be assigned to a host, which in 71% of cases was *Ralstonia* (summarized in Table S1, column “host prediction”).

Aeolian transport of viruses over polar environments, especially attached to snowflakes, has been barely investigated to date. Former investigations, mainly conducted at low latitudes, demonstrated intercontinental transport of microbes by winds (58), and that highly identical phages can be found in distantly related areas and in various ecosystems around the globe (59–62). C. M. Bellas et al. (63) recently reported the presence of near-identical phage genomes being spread by up to 4000 km in cryoconite holes of Svalbard, Greenland, and the Alps. Our data show that despite the isolation of the Antarctic continent, and under no consideration of anthropogenic dispersal (64), bacteria and phage distribution via snow over extensive distances across Antarctica is possible. Man-made dispersal seems unlikely for our samples, due to the relatedness of Antarctic phages to phages from environmental sources according to database hits. We cannot be certain about the nature of the PGFs (prophage or lytic phage) by metagenomic predictions alone. While the *Inoviridae* fraction likely occurs as prophages or episomal forms, detection of PGFs assigned to lytic categories by VIBRANT and VirSorter (category 1-3) suggests that host-associated, lytic phages captured at the adsorption or infection stages were present as well. The degree of uncertainty about the category of a virus presumably results from many fragmented scaffolds with relatively low coverages, which were however sufficient to identify shared elements of viruses between stations. Conclusions about the presence/absence of hallmark genes should be drawn from more complete datasets based on greater sequencing depth of samples containing more biomass. We further commend experiments that involve cultivation attempts and sequencing of metaviromes (free phages) to reveal the extent of lysogenic or chronic compared to lytic infection styles in Antarctic snow.

Snow PGFs carried identical SNPs with those from the seawater metagenome from the WAP located 4710 km and 5338 km apart from Druzhnaja and Mirnii, respectively (Figure 3C), implying long distance transport. This is further supported by the detection of their hosts, *Ralstonia* and *J. lividum,* in the WAP. From seawater, a transmission route via sea spray aerosols to snow that is blown over ice surfaces (65) can be assumed. Aerosolization of bacteria from the sea surface is highly taxon-specific but seems to work well for *R. pickettii* (66) and also viruses (67). Alternatively, a transport via aerosols to clouds and precipitating snow is possible. In the latter scenario, bioaerosols including microbial cells would act as ice nucleation particles (68), transferring microbes to ice clouds where they might induce their own precipitation (69, 70). The abundance data support the isolation of Progress-derived PGFs and their potential host *F. micromati* and point towards a decreasing gradient of abundances from Druzhnaja to Mirnii to the WAP for many PGFs (Table S1). Thus, general dispersal patterns of microbes across Antarctica seem governed by westward drift and are probably mediated by the prevailing Southern Hemisphere westerly winds (71). On short spatial-temporal scales, dispersal seems more complex and is probably shaped by multifactorial dependencies such as the different potential of a species to become airborne (66), meteorological conditions or the local geography. For instance, while Druzhnaja is located ∼50 km into the continent, the stations Mirnii and Progress are near the coast and more exposed to the sea in summer when ice breaks occur.

We assume that the transferability of the use of PGFs to study microbial dispersal in space analogues such as the Antarctic continent is likely applicable to other celestial bodies like Mars. Three celestial bodies in our solar system (Mars, Europa, and Enceladus) have environmental conditions that could favor microbial life (reviewed by M. G. Netea et al. (72)). Most microbial isolates (85-95%) obtained from spacecraft and assembly facilities are associated with humans (73). Since *Ralstonia* spp. can be human and plant pathogens (42, 74) and are able to thrive under harsh and oligotrophic conditions (7), they might contribute to the transmission of viruses to extraterrestrial environments, particularly via manned missions.

We found evidence for viral signatures associated with two *R. pickettii* strain draft genomes previously obtained from space equipment of a spacecraft assembly clean room and from water systems of the ISS. This result in conjunction with the result that temperate *Ralstonia* phages (and other viruses) can undergo long-range dispersal in association with their hosts across the extraterrestrial analogue Antarctica suggest that contaminations of space equipment with particularly persistent microbes such as *Ralstonia* should receive more focus during microbiological monitoring in the framework of Planetary Protection. However, the field of astrovirology has so far generally found little attention (75). The contribution of lysogenic and episomal phage (‘hidden hitchhikers’) to overall viral loads on spacecraft and associated equipment has been overlooked despite early work reporting on alterations in prophage induction during spaceflight (76–78), tobacco mosaic virus to survive space flight equivalent proton irradiation (79), the occurrence of phages and human-related circoviruses in clean rooms (80) and inoviruses on the ISS (81, 82).

Planetary Protection aims to prevent the spread of biological contaminants (forward contamination) to space shuttles and stations as well as extraterrestrial environments of the solar system. However, this policy largely ignores the potential of escaped biological contaminants to heavily disperse on foreign celestial bodies once being released, for instance after crash landings as happened for the Schiaparelli module of the ExoMars program in 2016 (83). Our results show that host-associated PGFs are not only suitable indicators for tracking long-distance dispersal in space analogues but also demonstrate that the release of contaminants that previously escaped Planetary Protection measures could spread far across extraterrestrial ecosystems and, in the worst-case scenario, confound future life detection missions.

## Material and Methods

### Metagenomic and genomic data processing

We analyzed publicly available metagenomic data sets, which correspond to Antarctic surface snow collected around three Russian stations (Druzhnaja, Mirnii, Progress) and are deposited at NCBI’s Sequence Read Archive (SRA) as Bioproject PRJNA674475 (Table S8) and MG-RAST under project accession mgp13052 including taxon abundance data. Snow sampling was conducted in December 2008 and 2009 as described previously (15); in brief, a sterile plastic scoop was used to sample ∼10 kg corresponding to a 2-3 cm layer of surface snow across several 1 m^2^ areas. Snow was melted over a period of 12 hours and concentrated using Pellicon tangential flow filters (Millipore, Burlington, MA, USA) to a final volume of ∼ 10 mL. DNA extraction for these metagenomes was carried out with the DNA Blood & Tissue kit (Qiagen, Hilden, Germany), whose protocol causes removal of free viruses, and resulted in DNA amounts of 170 – 490 ng. Sequencing libraries were prepared with MiSeq reagent kit v.2 (Illumina, USA), targeting mainly dsDNA viruses. Metagenomic data of a seawater sample from the WAP was also obtained from SRA (accession #SRR5591034). Raw shotgun sequencing reads of the three snow metagenomes and the WAP dataset were quality-trimmed using BBDuk (https://github.com/BioInfoTools/BBMap/blob/master/sh/bbduk.sh) from the BBTools package (84) and Sickle (85) resulting in read counts between 1.31 – 2.65 Mio. for the three snow samples. Assembly of reads was done using MetaSPAdes version 3.13 (86), and scaffolds <1 kbps were removed. Draft genome sequences of *R. pickettii* strains SSH4 and CW2 isolated from space equipment were obtained from Genbank accession #JFZG00000000 and #JFZH00000000. Strains SSH4 and CW2 were isolated pre-flight from the surface of the Mars Odyssey Orbiter during assembly and from a water sample taken in-flight from the ISS cooling system, respectively (9).

### Reconstruction of microbial genomes from metagenomes and CRISPR prediction

Binning of MAGs from the three stations Druzhnaja, Mirnii and Progress was performed using Emergent self-organizing maps (ESOMs, (87)), ABAWACA (https://github.com/CK7/abawaca) and MaxBin 2.0 (88). Aggregation of bins was performed using DASTool (89) and curation of bins was done in uBin (90), also delivering contamination and completeness scores (91). Read coverage of recovered MAGs was obtained from read mapping using Bowtie2 (92) in sensitive mode to the individual bins, followed by mismatch filtering (2% mismatch allowance, depending on read length). To investigate the dispersal of MAGs, mismatch criteria were set to 0% mismatches (100% similarity) and calcopo.rb (https://github.com/ProbstLab/viromics/tree/master/calcopo) was used to calculate the coverage per nucleotide (breadth) and the percentage of positions in the genome covered by reads. Only genomes with a least 70% breadth were considered present in the respective metagenome.

CRISPR arrays, which represent the prokaryotic adaptive immunity, were searched in host MAGs using CRISPRcasFinder and considering evidence level 3 or 4 (29). CRISPR loci consist of repeats interspaced with short DNA sequences (spacers) obtained from invading mobile genetic elements such as phages and thus provide a record of past infections. Since CRISPR arrays might get lost during the binning processes, e.g., due to fragmentation in assembly of strain variants, absence of CRISPR arrays in CRISPRcasFinder was further investigated by reconstructing CRISPR systems from raw reads using Crass (93) and BLASTing the obtained direct repeat (DR) sequences, which are phylogenetically well-conserved (94), against the NCBI non-redundant database (release 1^st^ March 2021) using BLASTn --short algorithm (95) with subsequent filtering at 80% similarity and e-value threshold of 10e-05. DR sequences were BLASTed against the MAGs and used to extract CRISPR spacers from reads using MetaCRAST with settings -d 3 -l 60 -c 0.99 -a 0.99 -r (96). Spacers were matched against PGFs as mentioned above for DRs. In most cases, BLASTing the DR sequences against the NR database did not reveal the host’s identity. Nevertheless, spacers derived from unknown hosts were considered, as their matches show that targeted PGFs represent true mobile genetic elements, and matches can be used to infer infection patterns between stations.

### H2PGF detection, host allocation and viral clustering

PGFs were identified from metagenome assemblies using a combination of bioinformatic tools, namely Virsorter v1 (33), VirFinder (97), CircMG (98), renamed to VRCA (https://github.com/alexcritschristoph/VRCA), VOGdb (version VOG93) (99), and Endmatcher (https://github.com/ProbstLab/viromics/tree/master/Endmatcher). The classification of predicted PGFs as “putative viruses” and “viruses” was done as previously described in the Supplementary Figure 3 of (100). VIBRANT v.1.2.1 (30) with default settings was used to find additional PGFs. No length cut-off was set (101), since many known *Ralstonia* phages and inoviruses have genome sizes <10 kb (36, 50), and because of the low biomass of Antarctic snow samples little PGFs compared to other ecosystems were expected. VirSorter and VIBRANT aided in (pro)phage detection in draft genomes of *R. pickettii* from space equipment. CheckV (32) was used to determine the type of identified PGF and completeness, and viralComplete (31) was applied to predict closely related phages. PGFs associated with MAGs were identified by grepping the scaffold ID on the bins. Pairwise comparisons of Antarctic PGFs and clustering was done using nucleic acid- and amino acid-based VICTOR (39). The resulting intergenomic distances were used to infer a balanced minimum evolution tree with branch support via FASTME including Subtree Pruning and Regrafting (SPR) postprocessing (102) using the distance formula D0. Branch support was inferred from 100 pseudo-bootstrap replicates each. Trees were rooted at the midpoint (103) and visualized with FigTree v1.4.4 (104). Taxon boundaries at the species and genus level were estimated with the OPTSIL program (105), using the recommended clustering thresholds (39) and an F value (fraction of links required for cluster fusion) of 0.5 (106).

Clustering of PGFs from Antarctic snow and space equipment was further substantiated via vConTACT2 v.0.9.19 (40, 107) in combination with the ProkaryoticViralRefSeq database (v94, (108)) followed by visualization of viral clusters (VC) in Cytoscape v. 3.8.2 (109). Virus-host matches were determined using the tool VirHostMatcher (34) with d_2_∗ oligonucleotide frequency dissimilarity measures for a k-mer length of 6. Viral genus clusters were determined using VIRIDIC (110). Venn diagrams were calculated using the VIB-ugent webtool (http://bioinformatics.psb.ugent.be/webtools/Venn/).

### Mapping of reads to assembled PGFs and nucleotide polymorphism analysis

We assume that presence of a PGF at two locations confirmed by read mapping represents dispersal. To determine if an assembled PGF from a single sample occurred in other metagenomes even if not being assembled, we performed read mapping following the previously published guidelines (101): that reads should map to a PGF with at least 90% identity (Bowtie2 settings as in (111)), and more than 75% of scaffold should have a coverage of at least 1x. To detect breadth of a PGF, we again used calcopo.rb (see above). Mean coverage of PGFs was calculated using calc_coverage_v3 (https://github.com/ProbstLab/uBin-helperscripts/blob/master/bin/04_01calc_coverage_v3.rb) and normalized to sequencing depth. Analysis of nucleotide polymorphisms was conducted for the 13 PGFs that underwent long range dispersal, i.e., PGF being present in the WAP sample and at least one of the snow metagenomes based on read mapping (Table S1). Variant analysis was performed in Geneious 11.1.5 (112) by applying default settings to the read-mappings generated as explained above.

### Gene prediction and annotations

Open reading frames on PGFs were detected using Prodigal in meta mode (113). Functional and taxonomic annotations of predicted proteins of PGFs were performed by DIAMOND searches with a e-value of 10e-05 (114) against FunTaxDB (90) and by using DRAM-v (115). For the full-length genome of the PGF p12D, annotations were improved using HHpred against PDB, Pfam, UniProt-SwissProt-viral and NCBI_Conserved Domains ((116, 117), https://toolkit.tuebingen.mpg.de/tools/hhpred) with a probability threshold of 70%. Sequences of PGFs were BLASTed against IMG/VR 2.0 (118) with an e-value cut-off of 10e-05 to find related phages from other metagenomic datasets.

### Synteny of *Ralstonia* phage and phylogenetic comparison of the *zot* gene

Synteny of known *Ralstonia* sp. phages from NCBI and all Antarctic PGFs that were identified herein and carried the zonular occludens toxin (*zot*, Pfam-ID: PF05707) was performed with tBLASTx comparisons using Easyfig v.2.2.5 (119) on .gbk files generated by Prokka (120). A phylogenetic tree for the MUSCLE-aligned (121) amino acid and nucleic acid sequences of *zot* was constructed using the FastTree (122) algorithm in Geneious 11.1.5 (112).

## Acknowledgements

We acknowledge Ken Dreger for server administration and maintenance as well as Cristina Moraru for sharing insights on virus taxonomy.

## Author Contribution statement

J.R. designed the study, wrote the manuscript, and carried out the analyses with input from T.L.V.B.; A.L. and K.S. generated the raw data and performed the sampling; A.J.P conceptualized the project, provided supervision, resources, and was involved in data interpretation; All authors edited drafts of the manuscript.

