## Supplementary material for "Host-associated phages disperse across the extraterrestrial analogue Antarctica"

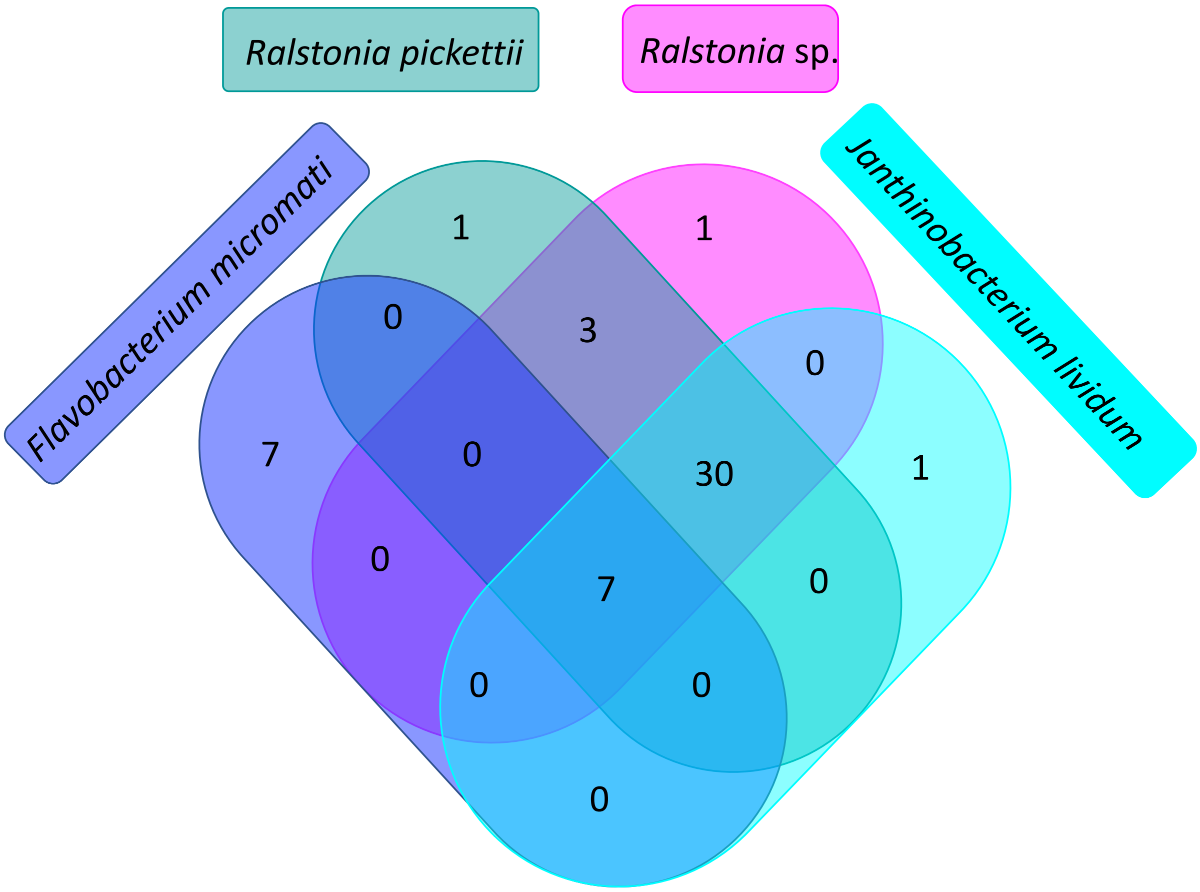


**Figure S1:** Venn-diagram showing overlapping phage-host matches of the 50 unique matching Antarctic PGFs as determined by VirHostMatcher (1).

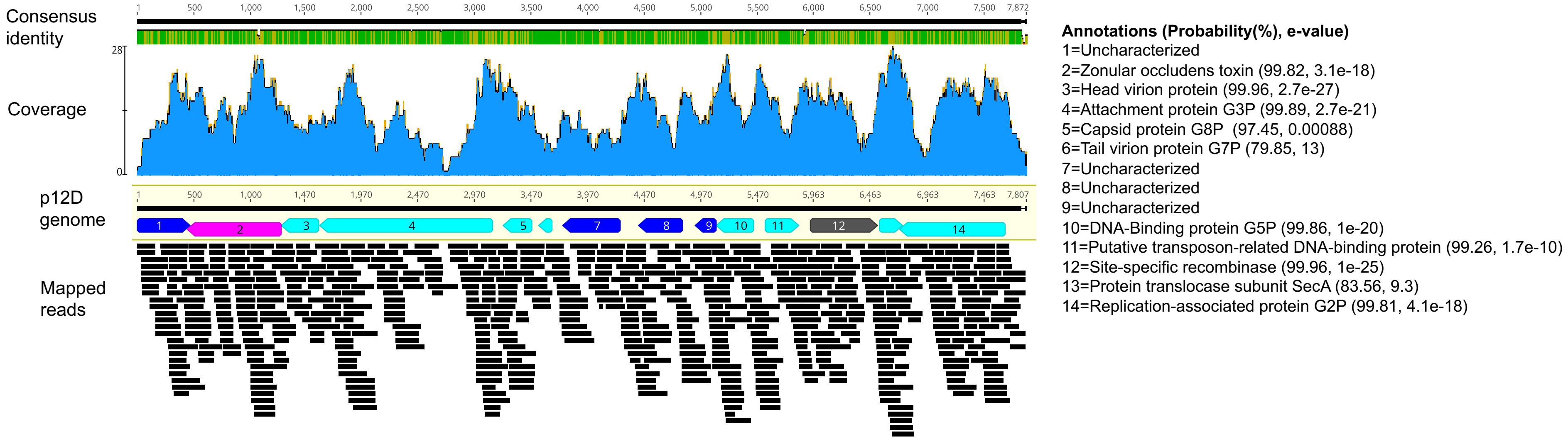


**Figure S2: Open reading frames and protein annotations of the phage genome of Antarcphage49_Dr_7823_circ**. Since this complete phage genome was found integrated in the chromosome of *Ralstonia* 12D, we propose the name *Ralstonia* phage p12D. Annotations of proteins were performed with HHpred (2, 3) against PDB, Pfam, UniProt-SwissProt-viral and NCBI_Conserved Domains using a probability threshold of 70%. Uncharacterized proteins are colored dark blue, the zot ORF in pink, the resolvase/recombinase in grey and remaining ORF in light blue. Reads from Druzhnaja were mapped against the genome using bowtie -sensitive mode (4). Genes that do not align by height overlap in their sequence. Visualization was performed in Geneious Prime version 11.1.5 (5).

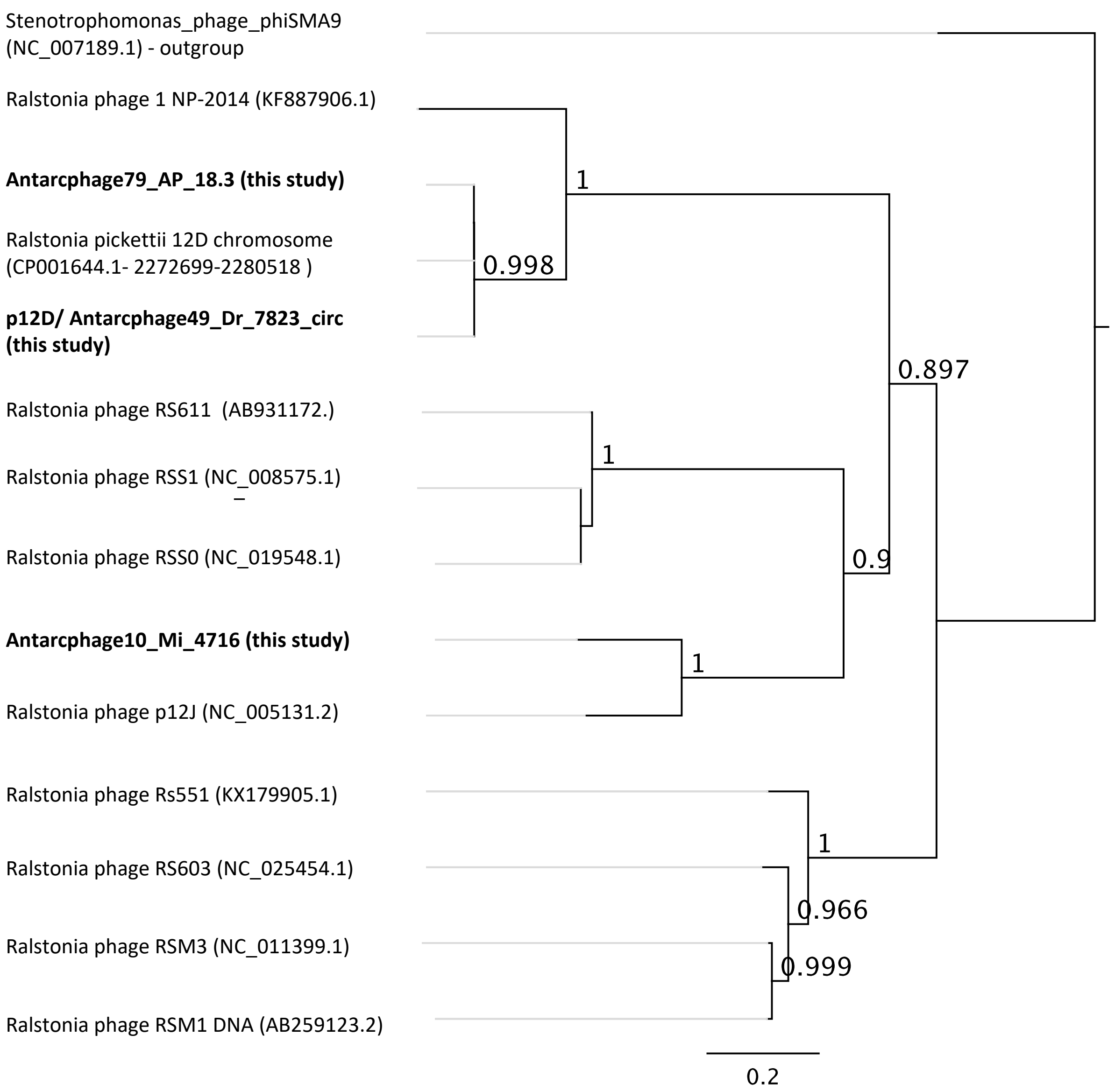


**Figure S3:** Phylogenetic tree constructed with FastTree 2.1.11 (6) in Geneious 11.1.5 under default settings for MUSCLE-aligned (7) nucleic acid sequence of the Zot protein of known *Ralstonia* phages and Antarctic phage genome fragments.

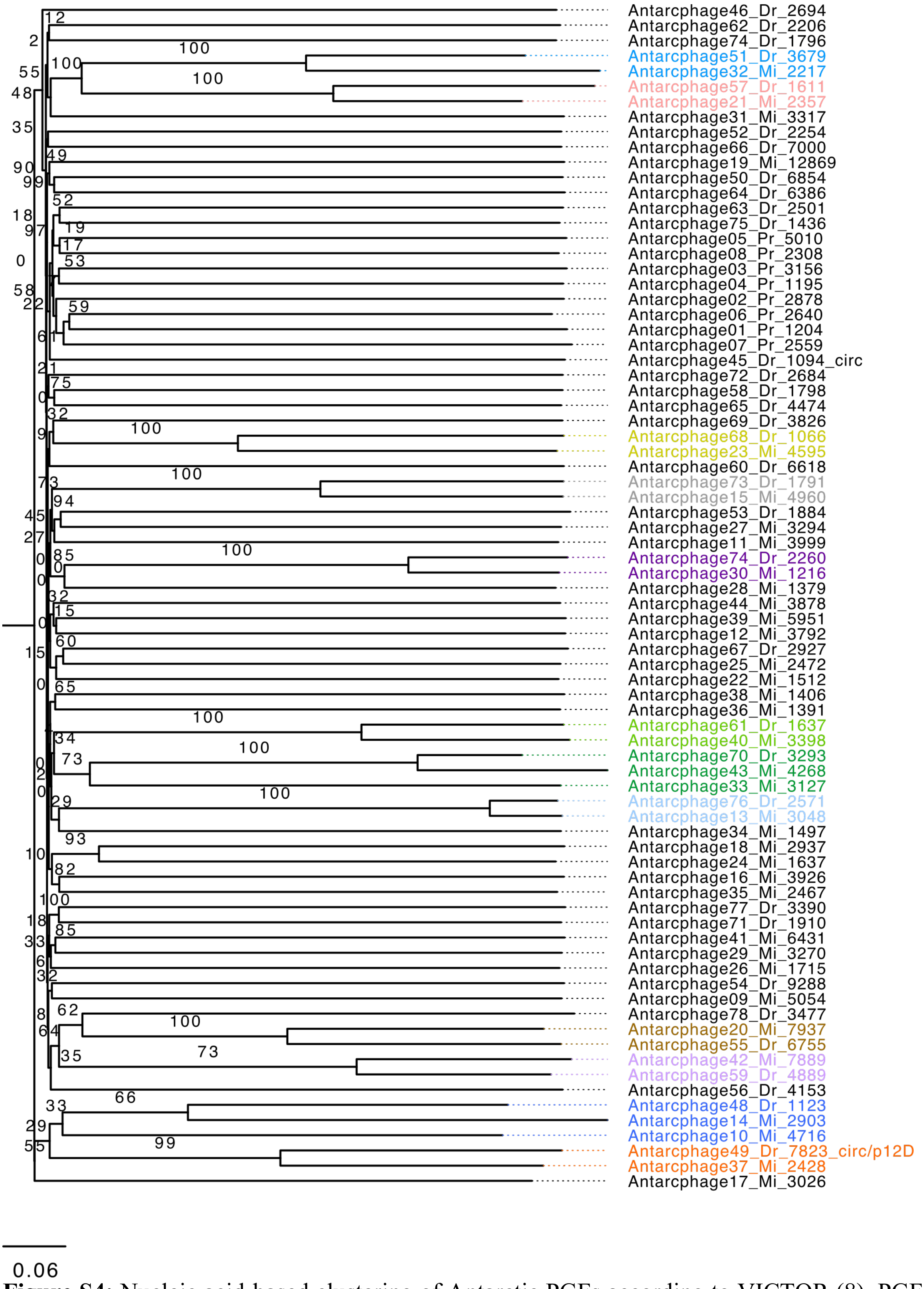


**Figure S4:** Nucleic acid-based clustering of Antarctic PGFs according to VICTOR (8). PGFs clustering on genus level (12 viral clusters) are color-coded. Branch support was inferred from 100 pseudo-bootstrap replicates each. Circ=circular, Dr=Druzhnaja, Mi=Mirnii (station), Pr=Progress (station). Visualization was performed with FigTree v.1.4.4. (9)

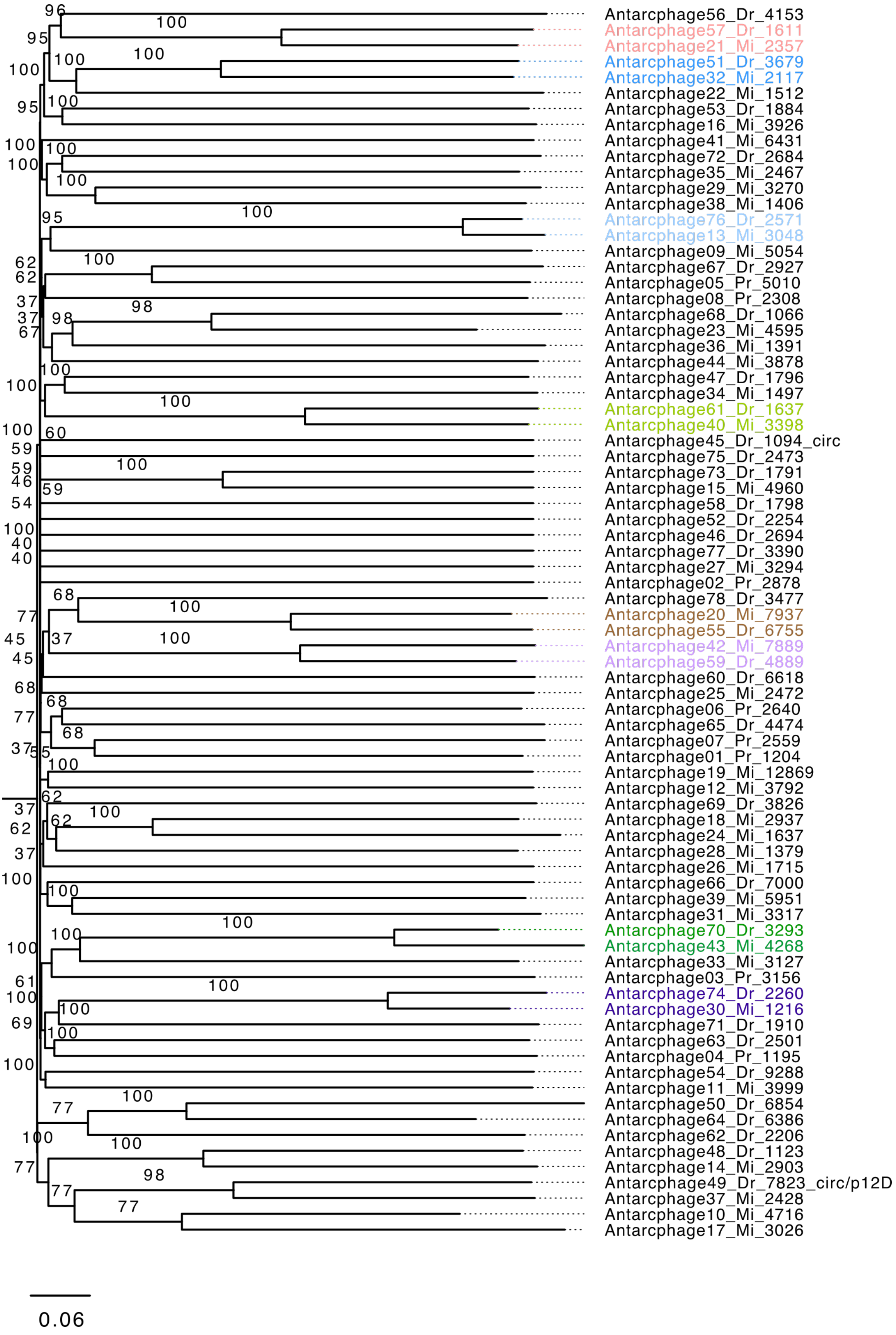


**Figure S5:** Amino acid-based clustering of Antarctic PGFs according to VICTOR (8). PGFs clustering on genus level (8 viral clusters) are color-coded. Branch support was inferred from

100 pseudo-bootstrap replicates each. Circ=circular, Dr=Druzhnaja, Mi=Mirnii (station), Pr=Progress (station). Visualization was performed with FigTree v.1.4.4. (9).
